# Analysis of the transcriptome and DNA methylome in response to acute and recurrent low glucose in human primary astrocytes

**DOI:** 10.1101/2020.07.07.191262

**Authors:** Paul G Weightman Potter, Sam Washer, Aaron R Jeffries, Janet E Holley, Nick J Gutowski, Emma Dempster, Craig Beall

**Author notes:** Co-corresponding authors: Emma Dempster, Institute of Biomedical and Clinical Sciences, University of Exeter Medical School, RILD Building, Barrack Road, Exeter, EX2 5DW, UK, 01392 408264, Craig Beall, Institute of Biomedical and Clinical Sciences, University of Exeter Medical School, RILD Building, Barrack Road, Exeter, EX2 5DW, UK, 01392 408209. Contributed equally (Paul G Weightman Potter, Sam Washer).

## Abstract

**Aims/hypothesis:** Recurrent hypoglycaemia (RH) is a major side-effect of intensive insulin therapy for people with diabetes. Changes in hypoglycaemia sensing by the brain contribute to the development of impaired counterregulatory responses to and awareness of hypoglycaemia. Little is known about the intrinsic changes in human astrocytes in response to acute and recurrent low glucose (RLG) exposure.

**Methods:** Human primary astrocytes (HPA) were exposed to zero, one, three or four bouts of low glucose (0.1 mmol/l) for three hours per day for four days to mimic RH. On the fourth day, DNA and RNA were collected. Differential gene expression and ontology analyses were performed using DESeq2 and GOseq respectively. DNA methylation was assessed using the Infinium MethylationEPIC BeadChip platform.

**Results:** 24 differentially expressed genes (DEGs) were detected (after correction for multiple comparisons). One bout of low glucose exposure had the largest effect on gene expression. Pathway analyses revealed that endoplasmic-reticulum (ER) stress-related genes such as *HSPA5*, *XBP1*, and *MANF*, involved in the unfolded protein response (UPR), were all significantly increased following LG exposure, which was diminished following RLG. There was little correlation between differentially methylated positions and changes in gene expression yet the number of bouts of LG exposure produced distinct methylation signatures.

**Conclusions/interpretation:** These data suggest that exposure of human astrocytes to transient LG triggers activation of genes involved in the UPR linked to endoplasmic reticulum (ER) stress. Following RLG, the activation of UPR related genes was diminished, suggesting attenuated ER stress. This may be mediated by metabolic adaptations to better preserve intracellular and/or ER ATP levels, but this requires further investigation.

## INTRODUCTION

Iatrogenic hypoglycaemia is a limiting factor to optimal glycaemic control in people with type 1 diabetes (T1DM; [1]). Recurrent hypoglycaemia (RH) leads to an impaired counterregulatory response (CRR) to restore blood glucose levels, at least in part mediated by the central detection of hypoglycaemia.

Astrocytes are glial cells with important roles in regulating neurotransmission, immune support, memory formation, long-term potentiation, and metabolic support to neurons. Astrocytes in the nucleus of the tractus solitarius (NTS) in the hindbrain increase intracellular Ca^2+^ levels in response to gluoprivation [2] and astrocytic glucose transporter GLUT2 is required for hypoglycaemia counterregulation [3]. Astrocytic glutamate uptake is impaired following RH [4], contributing to counterregulatory failure, yet little is known about the intrinsic changes within astrocytes, especially human astrocytes, following recurrent low glucose (RLG). In this study, we used both RNA sequencing and an epigenome-wide association study (EWAS) of DNA methylation (DNAm) to examine for the first time, changes to the human astrocyte transciptome and methylome following acute and recurrent low glucose exposure.

## RESEARCH DESIGN AND METHODS

### Astrocyte isolation and cell culture

HPA cells were isolated from post-mortem sub-ventricular deep white matter following consent from next-of-kin, and with ethical approval from the North and East Devon Research Ethics Committee as previously described [5]. The recurrent low glucose (RLG) model has been previously described (see ESM for details [6]). Samples were split for RNA extraction and DNA extraction, with a total of five and six replicates for RNA sequencing and DNA methylation studies, respectively. Cells were confirmed as mycoplasma free using the MycoAlert kit (Lonza, Slough, UK).

### RNA sequencing

Briefly, RNA was extracted and cDNA libraries generated. Sequencing reads were generated using the Illumina HiSeq 2500 and fastq sequence quality was checked using MultiQC before alignment to the human genome (Build GRCh38.p12) using STAR. Mapped reads were counted using the FeatureCounts function of the subread package. Differential gene expression was calculated using DESeq2 [7] using the Likelihood ratio test function to analyse all groups together followed by the Wald-test for pairwise analysis. Genes with a false discovery rate (FDR) ≤0.05 were considered differentially expressed. Functional gene ontology analysis was performed using GOSeq. See ESM for details. Raw RNAseq files are available at GEOLINK.

### DNA methylation analysis

DNA was extracted and DNA methylation (DNAm) examined using the Infinium MethylationEPIC BeadChip platform (Illumina Inc.; EPIC). See ESM for details of quality control and normalisation processes. 729727 probes remained after QC processes. The one-way analysis of variance (ANOVA) test was used to test for differentially methylated sites associated across the three groups: LG, aRLG, RLG compared to control. To determine which group was driving the association behind the significant ANOVA results, the *T* statistics for control versus each of the three groups were extracted from the regression model.

## RESULTS

### Low glucose-induced changes in gene expression in human astrocytes

In HPA cells, expression of 1240 genes were significantly (*p*<0.05) altered in response to glucose variation; 24 of which were significantly differentially expressed (DE) after FDR correction (adjusted *p*<0.05; Fig. 1A). Volcano plots displaying the pairwise comparisons of each treatment group versus control shows that LG (Fig. 1Ai) produced the largest effect on gene expression, whereas changes induced by aRLG (Aii) and RLG (Aiii) were more modest. LG and RLG shared similar DE patterns (Fig. 1B) and importantly *TXNIP*, regulated by glucose [8], was significantly downregulated in both LG (log2 fold-change −2.16, *p*=1.09E-5) and RLG (log2 fold-change −1.46, *p*=2.91E-3; Fig. 1C). Of the other DE genes there was a predominance of genes related to endoplasmic reticulum (ER)-stress. X-box binding protein 1 (*XBP1*; log2 fold-change 0.28, *p*=1.56E-4; Fig. 1D), heat shock protein family A member 5 (*HSPA5*; log2 fold-change 0.34, *p*=3.55E-6; Fig. 1E), and mesencephalic astrocyte-derived neurotrophic factor (*MANF*; log2 fold-change 0.41, *p*=7.55E-6; Fig. 1F) showed increased expression following LG exposure which was blunted following RLG. Similarly, mitochondrially encoded NADH:ubiquinone oxidoreductase core, subunit 4 and subunit 4L (*ND4* and *ND4L*) had increased gene expression in acute LG (ND4 log2 fold-change 0.37, *p*=3.5E-6; ND4L log2 fold-change 0.47, *p*=5.75E-7) and a diminished, but still significant increase following RLG (Fig. 1G,H). Pathway analysis of the DE genes identified seven gene ontology (GO) terms that were significantly altered after correction for multiple comparisons, which were related to the unfolded protein response (UPR) and ER-stress (ESM Table 1).

**Figure 1.**
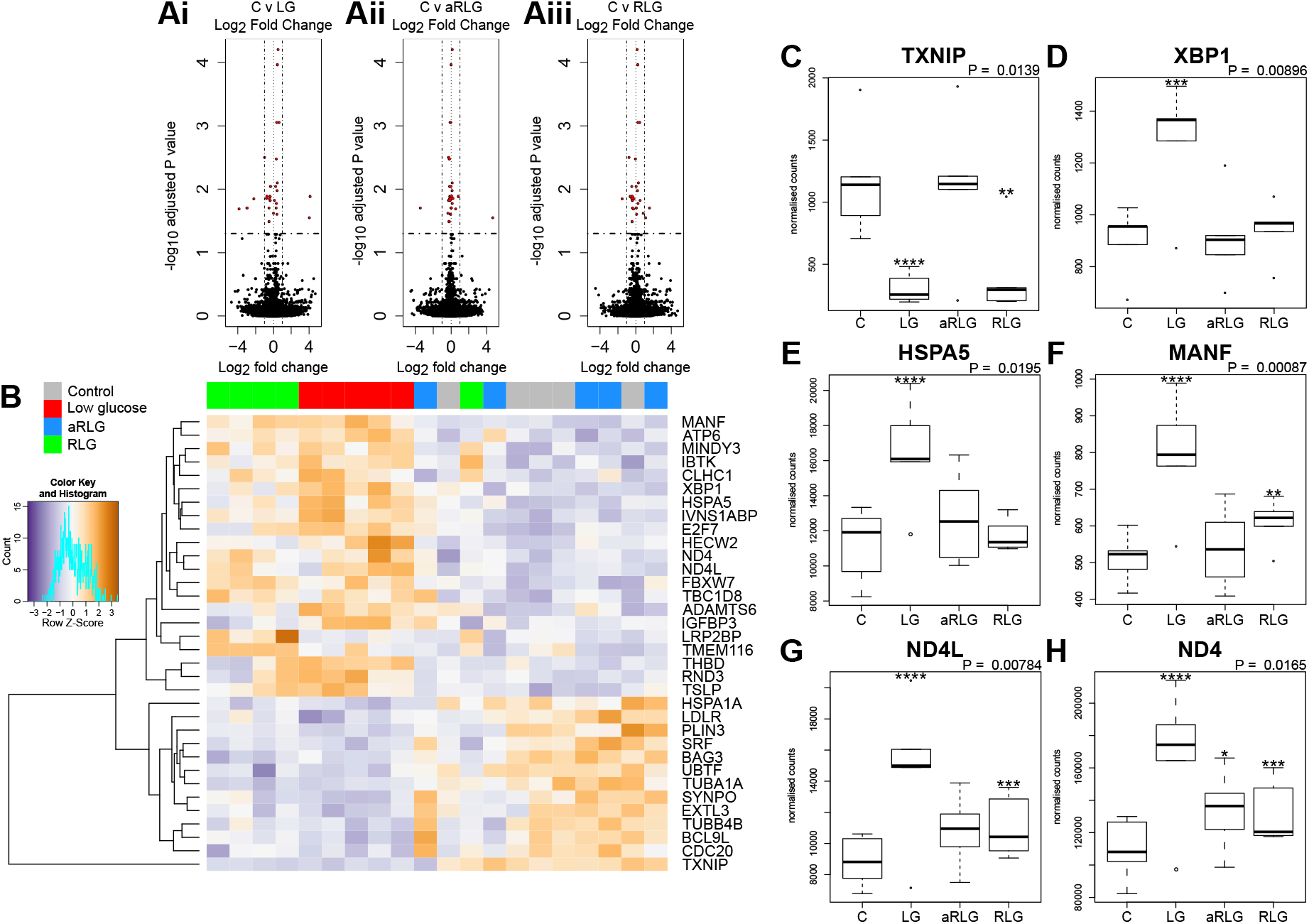
Summary of the RNAseq results. Volcano plots on the pairwise differential expression analysis between control cells (C) versus (**Ai**) low glucose (LG), (**Aii**) antecedent RLG, and (**Aiii**) recurrent low glucose (RLG), the red points on the plots represent genes padj<0.05. **B.** Heatmap of hierarchical clustering of LRT analysis FDR≤0.1 indicates differentially expressed genes (rows) between the four groups (padj<0.1). Orange indicates up-regulation and blue indicates down-regulation. The LG and RLG groups cluster together. TXNIP (**C**), XBP1 (**D**), HSPA5 (**E**), MANF (**F**), ND4L (**G**), ND4 (**H**) expression profiles, selected for their functional relevance to hypoglycaemia (*p*-value is the adjusted result of the likelihood ratio test). \**p*<0.05, \*\**p*<0.01, \*\*\**p*<0.001. Error bars represent standard deviation. N=5.

### LG and RLG produce distinct DNA methylation profiles

Our analyses did not identify any differential methylated positions (DMP) that reached genome-wide significance for DNA methylation association analyses (Fig. 2Ai-iii; *p*<9.42×10^−8^; [9]). However, 65 probes reached nominal significance of *p*<0.0001. Hierarchical clustering of these top probes showed four distinct groups that matched with the four experimental conditions suggesting a DNA methylation profile specific to each condition (Fig. 2B). Of the differentially methylated CpG sites, several were related to energy or ion homeostasis. *SLC19A3* (cg07417745, *p*=5.16E-7, βΔ= 0.23), encoding the thiamine transporter was hypermethylated after LG showing a linear relationship with the number of bouts of LG exposure (Fig. 2C). Similarly, methylation of the *GRID1* gene, encoding the ionotropic glutamate receptor δ1 (cg16777181) was hypermethylated following LG exposure (*p*=1.90E-3, βΔ=0.18) and this remained elevated following RLG (Fig. 2D). In contrast, cg1102254 (*NIPA1*, *p*=2.65E-6, βΔ= −0.02), cg11692715 (*SLC8B1*; *p*=1.61E-5, βΔ= −0.16) and cg22467827 (*CLHC1*, *p*=4.28E-4, βΔ= −0.03), which encode a Mg^2+^ transporter [10], a Na^+^/Ca^2+^ antiporter, and clathrin heavy chain linker domain containing 1 respectively, were hypomethylated following RLG (Fig. 2E-G). The probe cg22467827 (annotated to the gene *CLHC1*) was also differentially expressed (log2 fold-change 0.80, *p*=1.03E-4) in relation to RLG (Fig. 2H). The two datasets (RNAseq and EPIC) were integrated resulting in 28 DE genes that overlapped with 31 differentially methylation positions (Fig. 2I).

**Figure 2.**
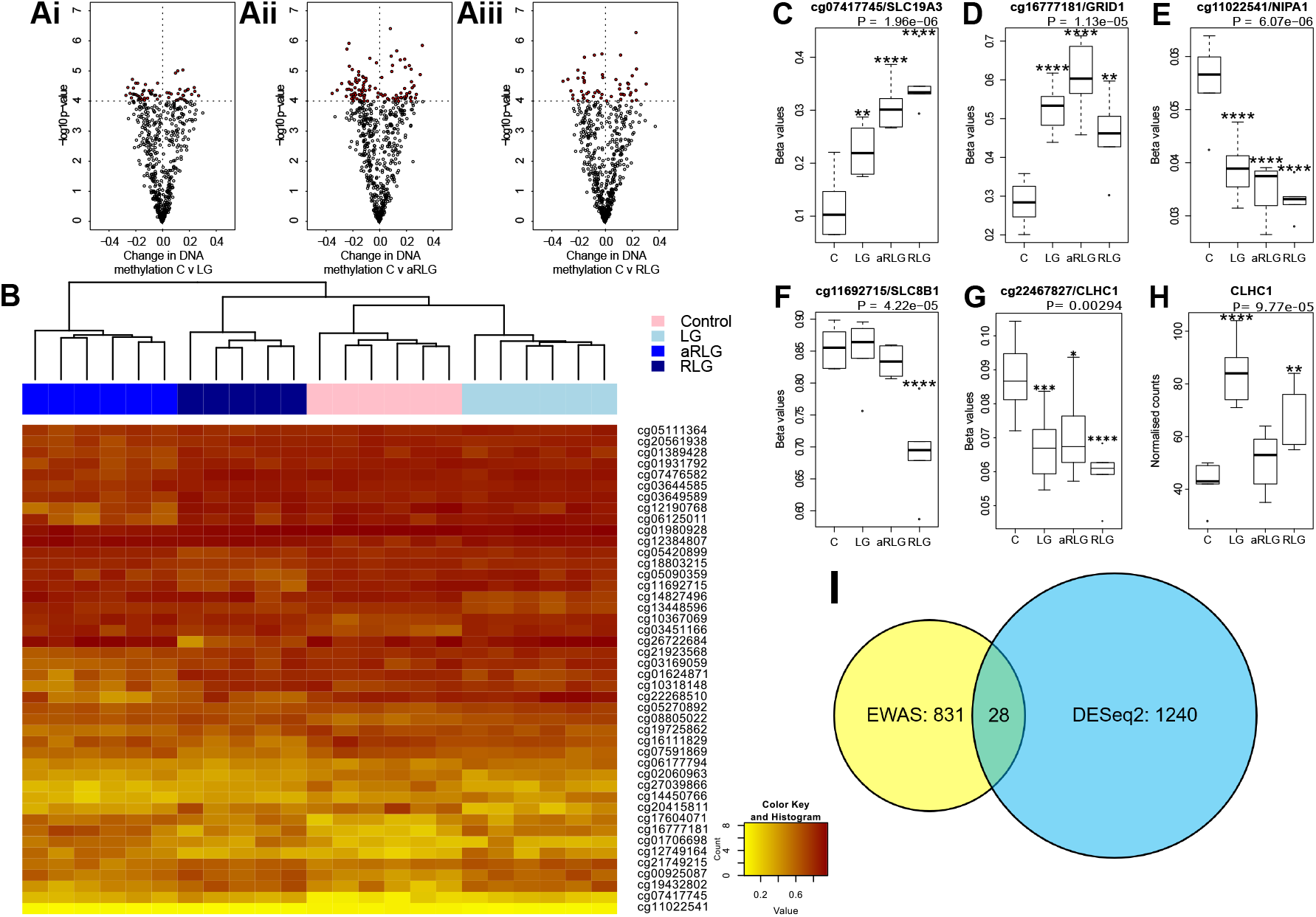
Summary of DNA methylation. **A.** the most differentially methylated genes (ANOVA p≤0.001) in pairwise comparison (red points are *p*<0.0001) between control treated HPA cells (C) versus (**Ai**) low glucose (LG), (**Aii**) antecedent recurrent low glucose (aRLG), and (**Aiii**) recurrent low glucose (RLG). **B**. Heatmap of hierarchical clustering using probes ANOVA *p*<0.001 indicates differentially methylated cg sites (rows) between the four groups. Orange indicates hypermethylation and yellow indicates hypomethylation. Box plots of some of the most differentially methylated cg sites labelled by their associated gene, selected for functional importance **C**, cg07417745/SLC19A3, **D**, cg16777181/GRID1, **E**, cg11022541/NIPA1, **F**, cg11692715/SLC8B1 **G**, cg22467827/CLHC1 (*p*-value is the adjusted result of the ANOVA). **H**, CLCH1 gene expression increases. Error bars represent standard deviation **I**, Venn diagram of differentially methylated cg sites in yellow and differentially expressed genes (blue) and overlap between the two data sets, 28 genes. Error bars represent standard deviation. \**p*<0.05, \*\**p*<0.01, \*\*\**p*<0.001, \*\*\*\**p*<0.0001. n=6 for methylation data and n=5 for gene expression changes.

## DISCUSSION

The central adaptations in response to RH that may mediate defective CRR require further investigation, with little known about how astrocytes respond or adapt to RH. We sought to examine changes in HPA gene expression and DNA methylation to determine which, if any, pathways were altered by acute and RLG exposure. DE and GO pathway analyses revealed that the major pathway altered by acute LG was the UPR. Protein folding within the ER requires hydrolysis of ATP (for review see [11]) and reductions in ATP content driven by energy stress increases protein misfolding to activate the UPR [12]. ER stress, via ATF6 promotes the production of XBP1 [13], which is spliced by IRE1α, to produce a potent transcriptional activator, XBP1s that increases HSPA5 [13] and MANF expression [14]). *MANF* is upregulated by UPR to inhibit cell proliferation and prevent ER-stress-related cell death [15, 16]. Interestingly, here expression of *XBP1*, *HSPA5*, and *MANF* were increased following a single bout of LG. Similar ER stress responses have been reported in pericytes [17], cardiac tissue [18], rat primary astrocytes [19] and primary hippocampal neurons [12] in response to LG. Following RLG, the increase in UPR-related gene expression was substantially diminished. Given that energy deficiency increases ER stress, it is plausible that acute LG exposure causes a marked increase in ER stress and that following successive bouts of LG, a concomitant metabolic adaptation, as previously reported [6], better preserves cellular ATP levels, thus attenuating (or delaying) subsequent LG-induced ER stress. In the data presented here, expression levels of *ND4L* and *ND4* following RLG remained elevated above control, suggesting a persistent adaptation. Whether this is mediated by better preservation of intracellular and intra-ER ATP levels remains to be determined.

The EWAS identified 65 DMPs associated with LG/RLG that reached nominal significance, while we did not identify any DMPs that reached the suggested array-wide significance (*p*<9.42×10^−8^). Hierarchical clustering revealed distinct patterns of DNA methylation across the four conditions. One of the most significant DMPs (cg07417745) is located in intron 1 of the *SLC19A3* gene, which encodes a thiamine transporter [20], and showed a linear relationship between increased DNA methylation and the number of LG exposures. Interestingly, expression of this gene has previously been found to be modulated by hyperglycaemic-like conditions [21]. Conversely, methylation of cg11022541 and cg11692715 located within the genes *NIPA1* and *SLC8B1* respectively, decreased following RLG. As these genes encode a Mg^2+^ transporter [10] and a Na^+^/Ca^+^ exchanger [22], this may indicate the energetic cost of ion handling within the cell, which requires further investigation. The main limitation in this study was that we were underpowered in the DNA methylation analyses as power analysis indicated we had 50% power to detect a difference of 10% in half of all the sites on the EPIC array. Moreover, the relationship between DNA methylation and gene expression is complex, with the direction of effect dictated by sequence context [23]. Furthermore, the annotation of DNAm sites to genes is purely based on proximity rather than empirical derived data [24], both these factors make inferences between DMPs and gene expression complicated. Despite these challenges we looked for overlapping genes between the datasets and identified 28 that were significantly altered (p <0.05) in both analyses. For example, *CLHC1* gene expression was significantly increased and a DMP (cg22467827) located in intron 1 was hypomethylated. This tentatively suggests that DNA methylation within the first intron may be mediating the upregulation of this gene in the response to LG glucose.

In summary, there are both shared and unique gene expression and DNA methylation profiles in human astrocytes following LG and RLG exposure. A single bout of LG exposure induced expression of genes associated with the UPR linked to ER stress. This response diminished after 4 bouts of LG exposure, suggesting an attenuated stress response. Taken together with previous observations that astrocytes adapt to RLG by increasing reliance on fatty acid oxidation to maintain intracellular ATP levels, activation of the UPR by glucose deprivation may be attenuated following RLG exposure.

## Acknowledgements

We thank the family of the donor for making this research possible. RNA library preparation and sequencing were performed by the Exeter Sequencing Service and Computational Core facilities at the University of Exeter.

## Contribution Statement

A.R.J, E.D. and C.B. conceived the project. P.G.W.P, S.W., A.R.J and E.D. contributed to data acquisition, analyses and writing of the manuscript. J.E.H and N.J.G obtained human post mortem tissue from which astrocytes were isolated, cultured and characterised as the stable human primary astrocyte population thus permitting the investigation of intrinsic changes in human astrocyte responses to be undertaken. C.B. wrote and edited the manuscript and accepts full responsibility for the work and/or conduct of the study, had access to the data and controlled the decision to publish.

---

**ESM figure 1.**
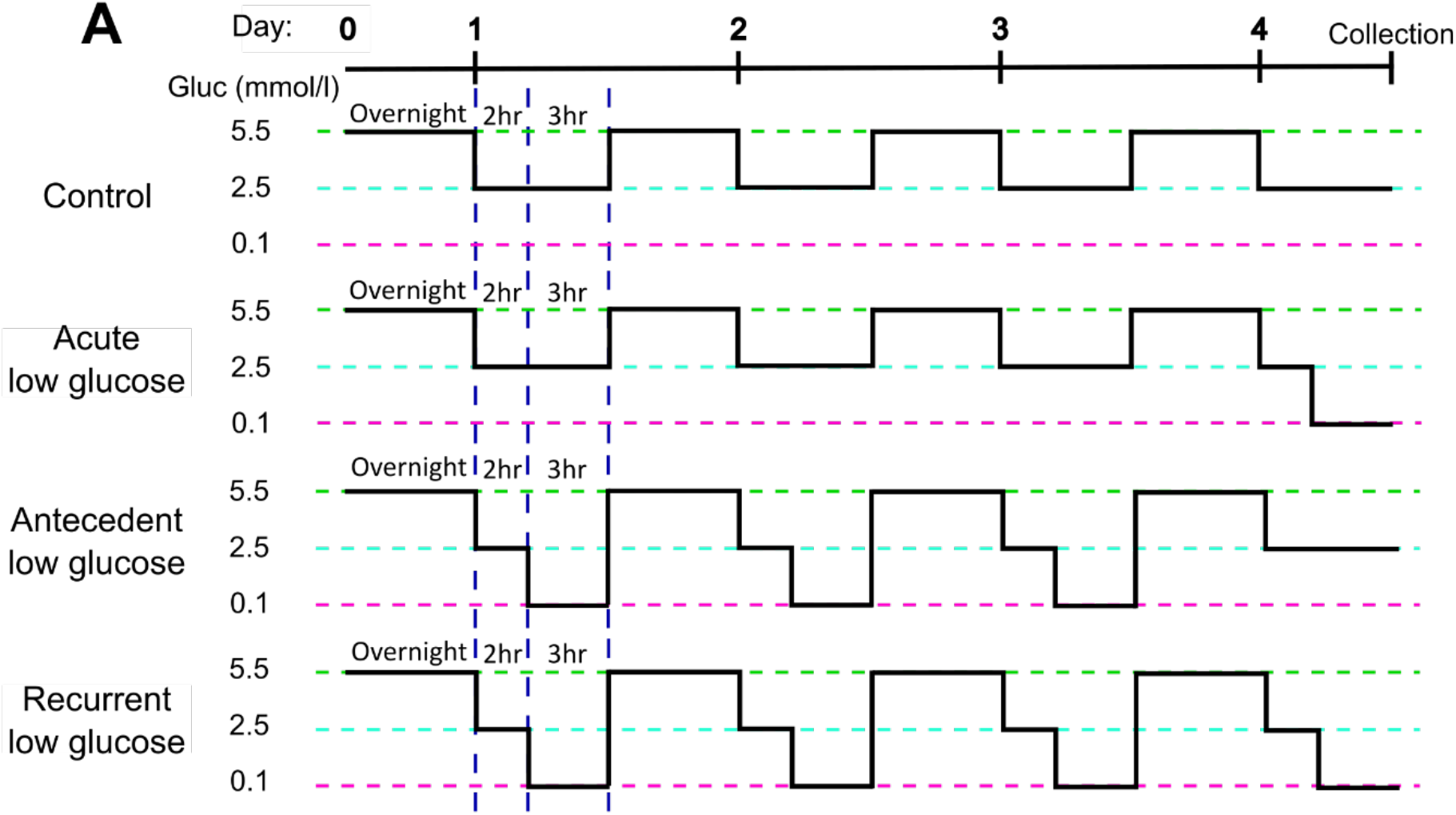
Schematic of the recurrent low glucose model. Human primary astrocytes were exposed to 0, 1, 3, or 4, three-hour long bouts of 0.1 mmol/l glucose; control (C), acute low glucose (LG), antecedent recurrent low glucose (aRLG), and recurrent low glucose (RLG) respectively. Each day cells were first incubated in 2.5 mmol/l glucose for 2 hours as a step down from overnight/stock media of 5.5 mmol/l glucose. Adapted from Weightman Potter *et al* 2019.

**ESM Table 1.**
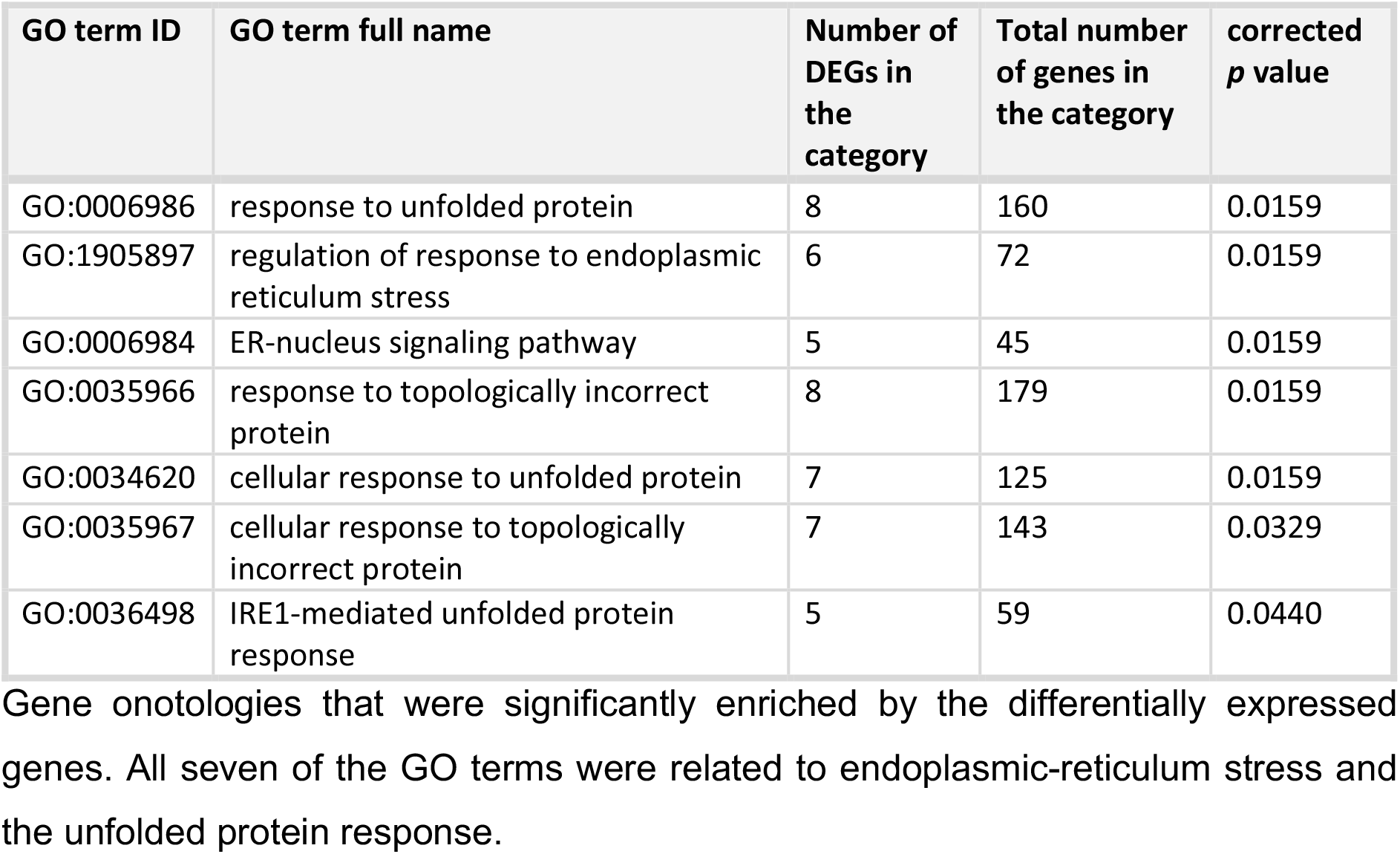
Glucose variation in human primary astrocytes significantly enriched gene ontologies related to endoplasmic-reticulum stress.

**ESM Table 2.**
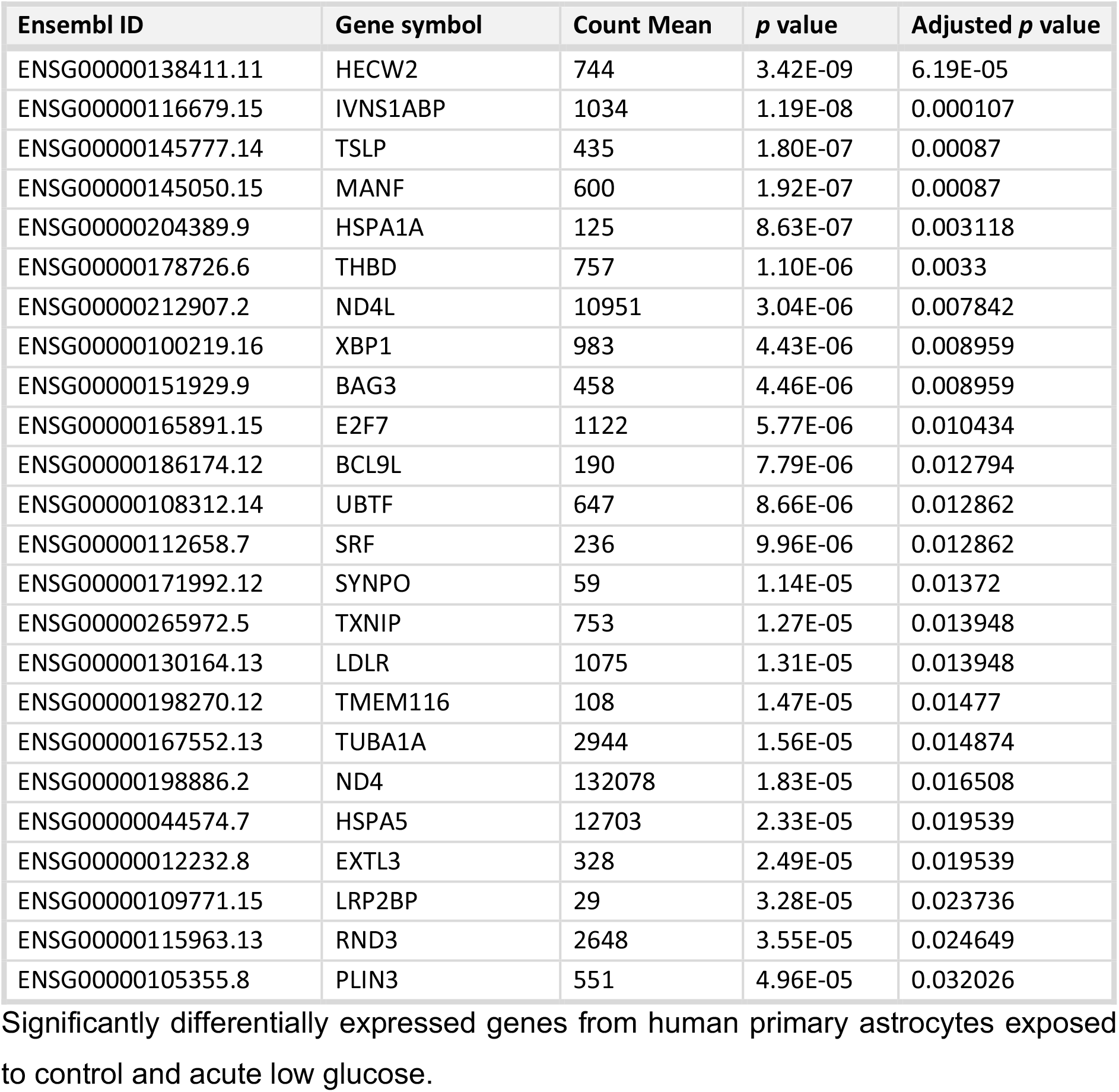
Glucose variation induces differentially expression of the following genes adjusted *p* value <0.05 in human primary astrocytes.

**ESM Table 3.**
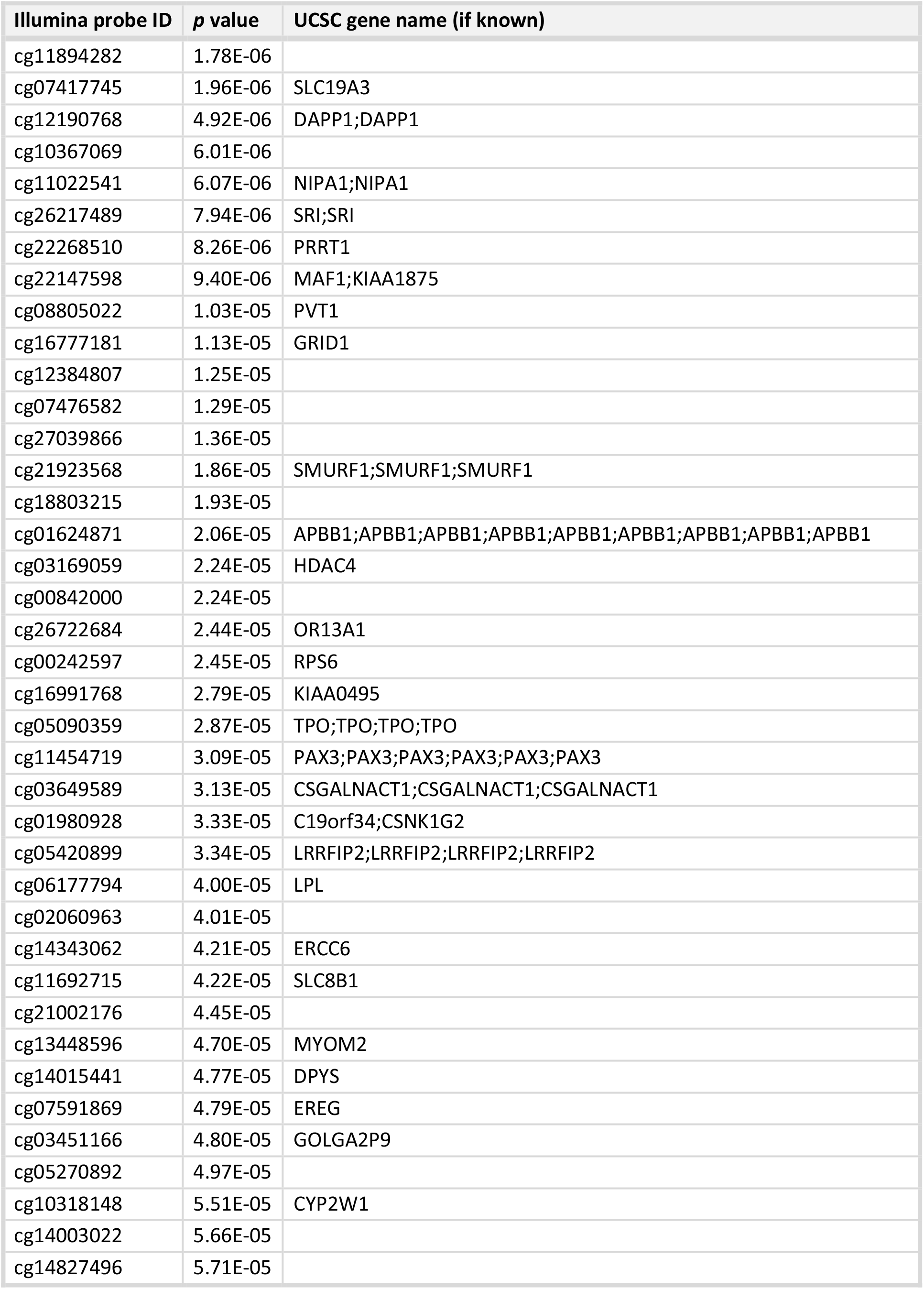
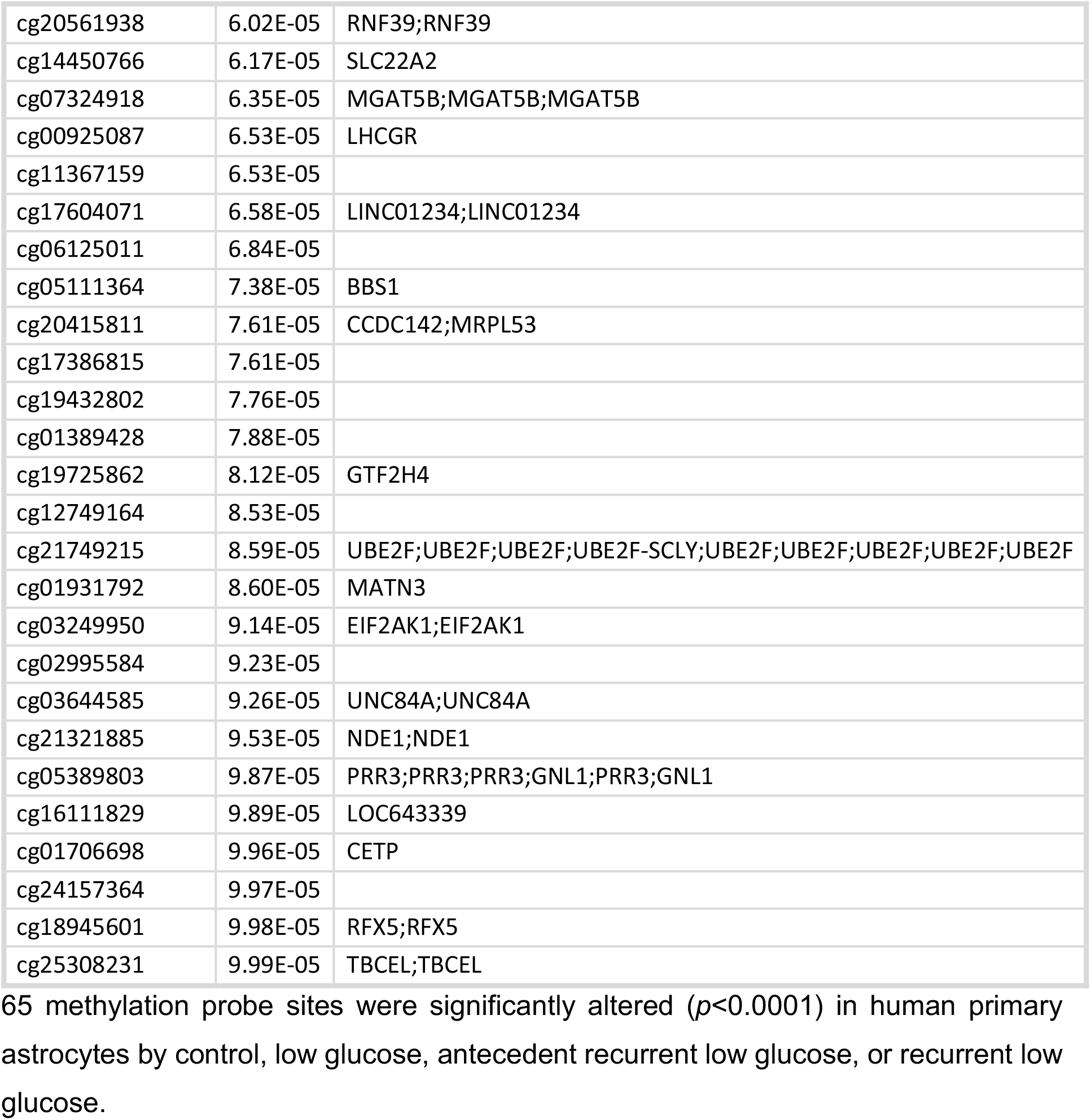
Glucose variation induces differentially methylation of the following cg sites *p* value <0.0001 in human primary astrocytes.

**ESM Table 4.**
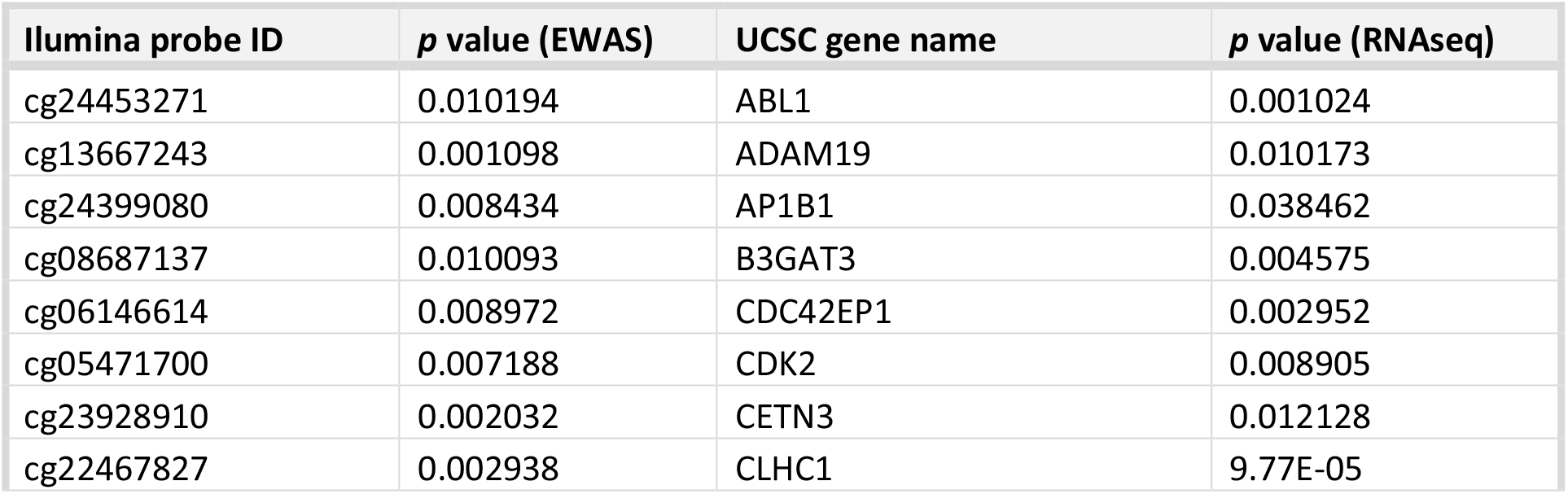
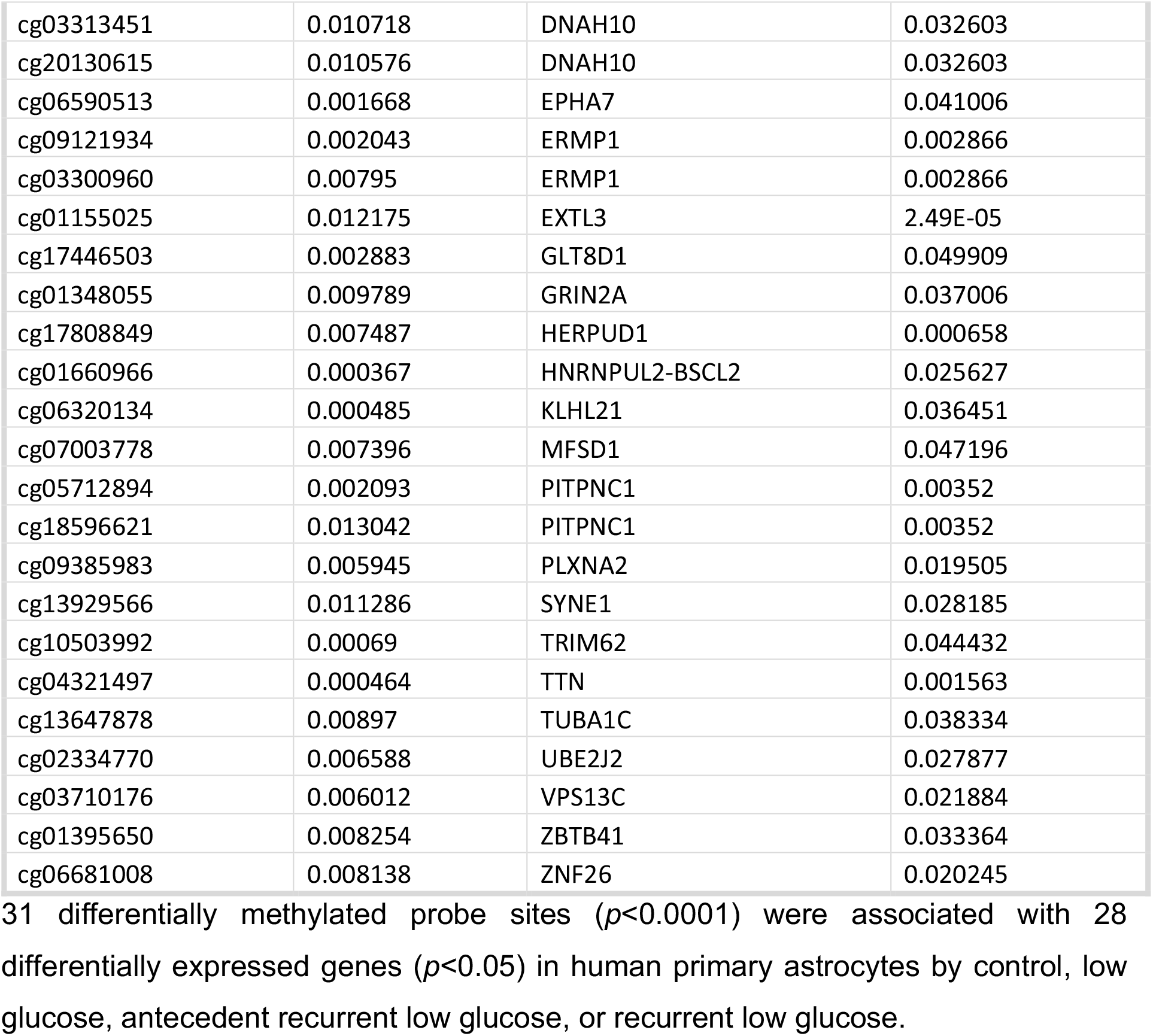
Glucose variation induces differential methylation of 31 probe sites associated with 28 differentially expressed genes in human primary astrocytes.

## Supplementary Methods

### Astrocyte isolation and cell culture

HPA cells were isolated from normal subventricular deep white matter blocks immediately post-mortem following consent from next-of-kin and with ethical approval from the North and East Devon Research Ethics Committee as previously described [5]. The recurrent low glucose (RLG) model has been previously described [6]. Each day cells were cultured in 2.5 mmol/L glucose-containing media for 2 hours before being changed for media containing 0.1 (low) or 2.5 (normal) mmol/L glucose for 3 hours. Overnight, cells were recovered in stock media containing 5.5 mmol/L glucose. This was repeated for four days. Control and low glucose (LG) treated cells had 2.5 mmol/L glucose for three days and on the fourth day the LG group received low glucose. The antecedent RLG (aRLG) and RLG groups had 0.1 mmol/L glucose for fourth days, except on the fourth day the aRLG groups had 2.5 mmol/L glucose. Samples were split for RNA extraction and DNA extraction, with a total of five and six replicates for RNA sequencing and DNA methylation studies, respectively. Cells were confirmed as mycoplasma free using the MycoAlert kit (Lonza, Slough, UK).

### RNA sequencing

Briefly, RNA was extracted using TRIzol and Direct-zol miniprep kit (Invitrogen, Carlsbad, CA, USA), according to manufacturers’ instructions. cDNA libraries were generated using the TruSeq DNA HT Library Preparation Kit (Illumina Inc., San Diego, CA, USA). Sequencing reads were generated using the Illumina HiSeq 2500 and fastq sequence quality was checked using MultiQC before alignment to the human genome (Build GRCh38.p12) using STAR. Mapped reads were counted using the FeatureCounts function of the subread package. Differential gene expression was calculated using DESeq2 using the Likelihood ratio test function to analyse all groups together followed by the Wald-test for pairwise analysis. Genes with a false discovery rate (FDR) ≤0.05 were considered differentially expressed. Functional gene ontology analysis was performed using GOSeq. Gene length was accounted for during GO analysis. Raw RNAseq files are available at GEOLINK.

### DNA methylation analysis

DNA was extracted using a modified phenol:chloroform protocol. DNA methylation was measured using the Infinium MethylationEPIC BeadChip platform (Illumina Inc.) (EPIC). 729727 probes remained after QC processes. The one-way analysis of variance (ANOVA) test was used to test for differentially methylated sites associated across the three groups: LG, aRLG, RLG compared to controls. To determine which group was driving the association behind the significant ANOVA results, the *T* statistics for controls versus each of the three groups were extracted from the regression model. Power was calculated using https://epigenetics.essex.ac.uk/shiny/EPICDNAmPowerCalcs/.

## References

1. Cryer, P.E., Hypoglycemia: still the limiting factor in the glycemic management of diabetes. Endocr Pract, 2008. 14(6): p. 750–6.

2. McDougal, D.H., G.E. Hermann, and R.C. Rogers, Astrocytes in the nucleus of the solitary tract are activated by low glucose or glucoprivation: evidence for glial involvement in glucose homeostasis. Frontiers in Neuroscience, 2013. 7.

3. Marty, N., M. Dallaporta, M. Foretz, M. Emery, D. Tarussio, I. Bady, C. Binnert, F. Beermann, and B. Thorens, Regulation of glucagon secretion by glucose transporter type 2 (glut2) and astrocyte-dependent glucose sensors. The Journal of clinical investigation, 2005. 115(12): p. 3545–3553.

4. Chowdhury, G.M.I., P. Wang, A. Ciardi, R. Mamillapalli, J. Johnson, W. Zhu, T. Eid, K. Behar, and O. Chan, Impaired Glutamatergic Neurotransmission in the VMH May Contribute to Defective Counterregulation in Recurrently Hypoglycemic Rats. Diabetes, 2017.

5. Holley, J.E., D. Gveric, J.L. Whatmore, and N.J. Gutowski, Tenascin C induces a quiescent phenotype in cultured adult human astrocytes. Glia, 2005. 52(1): p. 53–8.

6. Weightman Potter, P.G., J.M. Vlachaki Walker, J.L. Robb, J.K. Chilton, R. Williamson, A.D. Randall, K.L.J. Ellacott, and C. Beall, Basal fatty acid oxidation increases after recurrent low glucose in human primary astrocytes. Diabetologia, 2019. 62(1): p. 187–198.

7. Love, M.I., W. Huber, and S. Anders, Moderated estimation of fold change and dispersion for RNA-seq data with DESeq2. Genome Biol, 2014. 15(12): p. 550.

8. Parikh, H., E. Carlsson, W.A. Chutkow, L.E. Johansson, H. Storgaard, P. Poulsen, R. Saxena, C. Ladd, P.C. Schulze, M.J. Mazzini, C.B. Jensen, A. Krook, M. Björnholm, H. Tornqvist, J.R. Zierath, M. Ridderstråle, D. Altshuler, R.T. Lee, A. Vaag, L.C. Groop, and V.K. Mootha, TXNIP Regulates Peripheral Glucose Metabolism in Humans. PLOS Medicine, 2007. 4(5): p. e158.

9. Mansell, G., T.J. Gorrie-Stone, Y. Bao, M. Kumari, L.S. Schalkwyk, J. Mill, and E. Hannon, Guidance for DNA methylation studies: statistical insights from the Illumina EPIC array. BMC Genomics, 2019. 20(1): p. 366.

10. Goytain, A., R.M. Hines, A. El-Husseini, and G.A. Quamme, NIPA1(SPG6), the basis for autosomal dominant form of hereditary spastic paraplegia, encodes a functional Mg2+ transporter. J Biol Chem, 2007. 282(11): p. 8060–8.

11. Braakman, I. and D.N. Hebert, Protein folding in the endoplasmic reticulum. Cold Spring Harbor perspectives in biology, 2013. 5(5): p. a013201–a013201.

12. de la Cadena, S.G., K. Hernández-Fonseca, I. Camacho-Arroyo, and L. Massieu, Glucose deprivation induces reticulum stress by the PERK pathway and caspase-7- and calpain-mediated caspase-12 activation. Apoptosis, 2014. 19(3): p. 414–427.

13. Yoshida, H., T. Matsui, A. Yamamoto, T. Okada, and K. Mori, XBP1 mRNA is induced by ATF6 and spliced by IRE1 in response to ER stress to produce a highly active transcription factor. Cell, 2001. 107(7): p. 881–91.

14. Oh-Hashi, K., Y. Hirata, and K. Kiuchi, Transcriptional regulation of mouse mesencephalic astrocyte-derived neurotrophic factor in Neuro2a cells. Cell Mol Biol Lett, 2013. 18(3): p. 398–415.

15. Apostolou, A., Y. Shen, Y. Liang, J. Luo, and S. Fang, Armet, a UPR-upregulated protein, inhibits cell proliferation and ER stress-induced cell death. Experimental Cell Research, 2008. 314(13): p. 2454–2467.

16. Huang, J., C. Chen, H. Gu, C. Li, X. Fu, M. Jiang, H. Sun, J. Xu, J. Fang, and L. Jin, Mesencephalic astrocyte-derived neurotrophic factor reduces cell apoptosis via upregulating GRP78 in SH-SY5Y cells. Cell Biology International, 2016. 40(7): p. 803–811.

17. Ikesugi, K., M.L. Mulhern, C.J. Madson, K. Hosoya, T. Terasaki, P.F. Kador, and T. Shinohara, Induction of endoplasmic reticulum stress in retinal pericytes by glucose deprivation. Curr Eye Res, 2006. 31(11): p. 947–53.

18. Barnes, J.A. and I.W. Smoak, Glucose-regulated protein 78 (GRP78) is elevated in embryonic mouse heart and induced following hypoglycemic stress. Anat Embryol (Berl), 2000. 202(1): p. 67–74.

19. Lind, K.R., K.K. Ball, N.F. Cruz, and G.A. Dienel, The unfolded protein response to endoplasmic reticulum stress in cultured astrocytes and rat brain during experimental diabetes. Neurochem Int, 2013. 62(5): p. 784–95.

20. Eudy, J.D., O. Spiegelstein, R.C. Barber, B.J. Wlodarczyk, J. Talbot, and R.H. Finnell, Identification and Characterization of the Human and Mouse SLC19A3 Gene: A Novel Member of the Reduced Folate Family of Micronutrient Transporter Genes. Molecular Genetics and Metabolism, 2000. 71(4): p. 581–590.

21. Beltramo, E., A. Mazzeo, T. Lopatina, M. Trento, and M. Porta, Thiamine transporter 2 is involved in high glucose-induced damage and altered thiamine availability in cell models of diabetic retinopathy. Diabetes and Vascular Disease Research, 2020. 17(1): p. 1479164119878427.

22. Palty, R., W.F. Silverman, M. Hershfinkel, T. Caporale, S.L. Sensi, J. Parnis, C. Nolte, D. Fishman, V. Shoshan-Barmatz, S. Herrmann, D. Khananshvili, and I. Sekler, NCLX is an essential component of mitochondrial Na+/Ca2+ exchange. Proc Natl Acad Sci U S A, 2010. 107(1): p. 436–41.

23. Jones, P.A., Functions of DNA methylation: islands, start sites, gene bodies and beyond. Nature Reviews Genetics, 2012. 13(7): p. 484–492.

24. Hannon, E., T.J. Gorrie-Stone, M.C. Smart, J. Burrage, A. Hughes, Y. Bao, M. Kumari, L.C. Schalkwyk, and J. Mill, Leveraging DNA-Methylation Quantitative-Trait Loci to Characterize the Relationship between Methylomic Variation, Gene Expression, and Complex Traits. American journal of human genetics, 2018. 103(5): p. 654–665.

