## Supplementary figures for "Analysis of the transcriptome and DNA methylome in response to acute and recurrent low glucose in human primary astrocytes"

### ESM Fig. 1

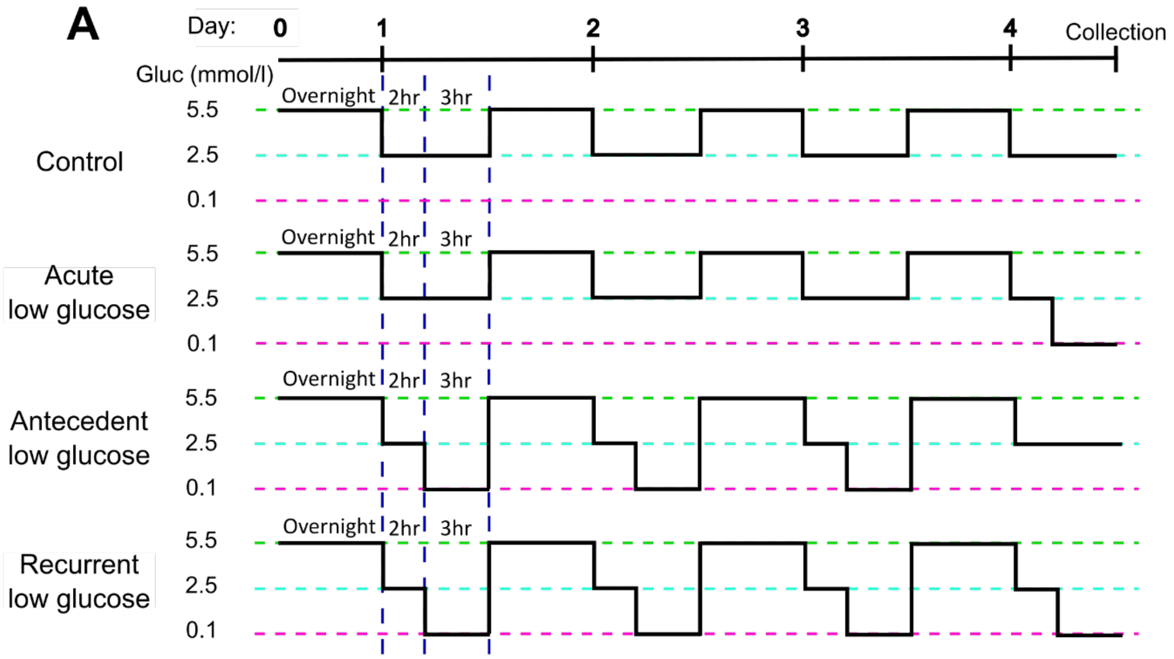


**ESM figure 1. Schematic of the recurrent low glucose model.** Human primary astrocytes were exposed to 0, 1, 3, or 4, three-hour long bouts of 0.1 mmol/l glucose; control (C), acute low glucose (LG), antecedent recurrent low glucose (aRLG), and recurrent low glucose (RLG) respectively. Each day cells were first incubated in 2.5 mmol/l glucose for 2 hours as a step down from overnight/stock media of 5.5 mmol/l glucose. Adapted from Weightman Potter *et al* 2019.

**ESM Table 1. Glucose variation in human primary astrocytes significantly enriched gene ontologies related to endoplasmic-reticulum stress**

| GO term ID | GO term full name | Number of DEGs in the category | Total number of genes in the category | corrected *p* value |
|---|---|---|---|---|
| GO:0006986 | response to unfolded protein | 8 | 160 | 0.0159 |
| GO:1905897 | regulation of response to endoplasmic reticulum stress | 6 | 72 | 0.0159 |
| GO:0006984 | ER-nucleus signaling pathway | 5 | 45 | 0.0159 |
| GO:0035966 | response to topologically incorrect protein | 8 | 179 | 0.0159 |
| GO:0034620 | cellular response to unfolded protein | 7 | 125 | 0.0159 |
| GO:0035967 | cellular response to topologically incorrect protein | 7 | 143 | 0.0329 |
| GO:0036498 | IRE1-mediated unfolded protein response | 5 | 59 | 0.0440 |

Gene onotologies that were significantly enriched by the differentially expressed genes. All seven of the GO terms were related to endoplasmic-reticulum stress and the unfolded protein response.

**ESM Table 2. Glucose variation induces differentially expression of the following genes adjusted *p* value <0.05 in human primary astrocytes**

| Ensembl ID | Gene symbol | Count Mean | *p* value | Adjusted *p* value |
|---|---|---|---|---|
| ENSG00000138411.11 | HECW2 | 744 | 3.42E-09 | 6.19E-05 |
| ENSG00000116679.15 | IVNS1ABP | 1034 | 1.19E-08 | 0.000107 |
| ENSG00000145777.14 | TSLP | 435 | 1.80E-07 | 0.00087 |
| ENSG00000145050.15 | MANF | 600 | 1.92E-07 | 0.00087 |
| ENSG00000204389.9 | HSPA1A | 125 | 8.63E-07 | 0.003118 |
| ENSG00000178726.6 | THBD | 757 | 1.10E-06 | 0.0033 |
| ENSG00000212907.2 | ND4L | 10951 | 3.04E-06 | 0.007842 |
| ENSG00000100219.16 | XBP1 | 983 | 4.43E-06 | 0.008959 |
| ENSG00000151929.9 | BAG3 | 458 | 4.46E-06 | 0.008959 |
| ENSG00000165891.15 | E2F7 | 1122 | 5.77E-06 | 0.010434 |
| ENSG00000186174.12 | BCL9L | 190 | 7.79E-06 | 0.012794 |
| ENSG00000108312.14 | UBTF | 647 | 8.66E-06 | 0.012862 |
| ENSG00000112658.7 | SRF | 236 | 9.96E-06 | 0.012862 |
| ENSG00000171992.12 | SYNPO | 59 | 1.14E-05 | 0.01372 |
| ENSG00000265972.5 | TXNIP | 753 | 1.27E-05 | 0.013948 |
| ENSG00000130164.13 | LDLR | 1075 | 1.31E-05 | 0.013948 |
| ENSG00000198270.12 | TMEM116 | 108 | 1.47E-05 | 0.01477 |
| ENSG00000167552.13 | TUBA1A | 2944 | 1.56E-05 | 0.014874 |
| ENSG00000198886.2 | ND4 | 132078 | 1.83E-05 | 0.016508 |
| ENSG00000044574.7 | HSPA5 | 12703 | 2.33E-05 | 0.019539 |
| ENSG00000012232.8 | EXTL3 | 328 | 2.49E-05 | 0.019539 |
| ENSG00000109771.15 | LRP2BP | 29 | 3.28E-05 | 0.023736 |
| ENSG00000115963.13 | RND3 | 2648 | 3.55E-05 | 0.024649 |
| ENSG00000105355.8 | PLIN3 | 551 | 4.96E-05 | 0.032026 |

Significantly differentially expressed genes from human primary astrocytes exposed to control and acute low glucose.

**ESM Table 3 Glucose variation induces differentially methylation of the following cg sites *p* value <0.0001 in human primary astrocytes**

| Illumina probe ID | *p* value | UCSC gene name (if known) |
|---|---|---|
| cg11894282 | 1.78E-06 | |
| cg07417745 | 1.96E-06 | SLC19A3 |
| cg12190768 | 4.92E-06 | DAPP1;DAPP1 |
| cg10367069 | 6.01E-06 | |
| cg11022541 | 6.07E-06 | NIPA1;NIPA1 |
| cg26217489 | 7.94E-06 | SRI;SRI |
| cg22268510 | 8.26E-06 | PRRT1 |
| cg22147598 | 9.40E-06 | MAF1;KIAA1875 |
| cg08805022 | 1.03E-05 | PVT1 |
| cg16777181 | 1.13E-05 | GRID1 |
| cg12384807 | 1.25E-05 | |
| cg07476582 | 1.29E-05 | |
| cg27039866 | 1.36E-05 | |
| cg21923568 | 1.86E-05 | SMURF1;SMURF1;SMURF1 |
| cg18803215 | 1.93E-05 | |
| cg01624871 | 2.06E-05 | APBB1;APBB1;APBB1;APBB1;APBB1;APBB1;APBB1;APBB1;APBB1 |
| cg03169059 | 2.24E-05 | HDAC4 |
| cg00842000 | 2.24E-05 | |
| cg26722684 | 2.44E-05 | OR13A1 |
| cg00242597 | 2.45E-05 | RPS6 |
| cg16991768 | 2.79E-05 | KIAA0495 |
| cg05090359 | 2.87E-05 | TPO;TPO;TPO;TPO |
| cg11454719 | 3.09E-05 | PAX3;PAX3;PAX3;PAX3;PAX3;PAX3 |
| cg03649589 | 3.13E-05 | CSGALNACT1;CSGALNACT1;CSGALNACT1 |
| cg01980928 | 3.33E-05 | C19orf34;CSNK1G2 |
| cg05420899 | 3.34E-05 | LRRFIP2;LRRFIP2;LRRFIP2;LRRFIP2 |
| cg06177794 | 4.00E-05 | LPL |
| cg02060963 | 4.01E-05 | |
| cg14343062 | 4.21E-05 | ERCC6 |
| cg11692715 | 4.22E-05 | SLC8B1 |
| cg21002176 | 4.45E-05 | |
| cg13448596 | 4.70E-05 | MYOM2 |
| cg14015441 | 4.77E-05 | DPYS |
| cg07591869 | 4.79E-05 | EREG |
| cg03451166 | 4.80E-05 | GOLGA2P9 |
| cg05270892 | 4.97E-05 | |
| cg10318148 | 5.51E-05 | CYP2W1 |
| cg14003022 | 5.66E-05 | |
| cg14827496 | 5.71E-05 | |

| | | |
|---|---|---|
| cg20561938 | 6.02E-05 | RNF39;RNF39 |
| cg14450766 | 6.17E-05 | SLC22A2 |
| cg07324918 | 6.35E-05 | MGAT5B;MGAT5B;MGAT5B |
| cg00925087 | 6.53E-05 | LHCGR |
| cg11367159 | 6.53E-05 | |
| cg17604071 | 6.58E-05 | LINC01234;LINC01234 |
| cg06125011 | 6.84E-05 | |
| cg05111364 | 7.38E-05 | BBS1 |
| cg20415811 | 7.61E-05 | CCDC142;MRPL53 |
| cg17386815 | 7.61E-05 | |
| cg19432802 | 7.76E-05 | |
| cg01389428 | 7.88E-05 | |
| cg19725862 | 8.12E-05 | GTF2H4 |
| cg12749164 | 8.53E-05 | |
| cg21749215 | 8.59E-05 | UBE2F;UBE2F;UBE2F;UBE2F-SCLY;UBE2F;UBE2F;UBE2F;UBE2F;UBE2F |
| cg01931792 | 8.60E-05 | MATN3 |
| cg03249950 | 9.14E-05 | EIF2AK1;EIF2AK1 |
| cg02995584 | 9.23E-05 | |
| cg03644585 | 9.26E-05 | UNC84A;UNC84A |
| cg21321885 | 9.53E-05 | NDE1;NDE1 |
| cg05389803 | 9.87E-05 | PRR3;PRR3;PRR3;GNL1;PRR3;GNL1 |
| cg16111829 | 9.89E-05 | LOC643339 |
| cg01706698 | 9.96E-05 | CETP |
| cg24157364 | 9.97E-05 | |
| cg18945601 | 9.98E-05 | RFX5;RFX5 |
| cg25308231 | 9.99E-05 | TBCEL;TBCEL |

65 methylation probe sites were significantly altered (*p*<0.0001) in human primary astrocytes by control, low glucose, antecedent recurrent low glucose, or recurrent low glucose.

**ESM Table 4 Glucose variation induces differential methylation of 31 probe sites associated with 28 differentially expressed genes in human primary astrocytes**

| Ilumina probe ID | *p* value (EWAS) | UCSC gene name | *p* value (RNAseq) |
|---|---|---|---|
| cg24453271 | 0.010194 | ABL1 | 0.001024 |
| cg13667243 | 0.001098 | ADAM19 | 0.010173 |
| cg24399080 | 0.008434 | AP1B1 | 0.038462 |
| cg08687137 | 0.010093 | B3GAT3 | 0.004575 |
| cg06146614 | 0.008972 | CDC42EP1 | 0.002952 |
| cg05471700 | 0.007188 | CDK2 | 0.008905 |
| cg23928910 | 0.002032 | CETN3 | 0.012128 |
| cg22467827 | 0.002938 | CLHC1 | 9.77E-05 |

| | | | |
|---|---|---|---|
| cg03313451 | 0.010718 | DNAH10 | 0.032603 |
| cg20130615 | 0.010576 | DNAH10 | 0.032603 |
| cg06590513 | 0.001668 | EPHA7 | 0.041006 |
| cg09121934 | 0.002043 | ERMP1 | 0.002866 |
| cg03300960 | 0.00795 | ERMP1 | 0.002866 |
| cg01155025 | 0.012175 | EXTL3 | 2.49E-05 |
| cg17446503 | 0.002883 | GLT8D1 | 0.049909 |
| cg01348055 | 0.009789 | GRIN2A | 0.037006 |
| cg17808849 | 0.007487 | HERPUD1 | 0.000658 |
| cg01660966 | 0.000367 | HNRNPUL2-BSCL2 | 0.025627 |
| cg06320134 | 0.000485 | KLHL21 | 0.036451 |
| cg07003778 | 0.007396 | MFSD1 | 0.047196 |
| cg05712894 | 0.002093 | PITPNC1 | 0.00352 |
| cg18596621 | 0.013042 | PITPNC1 | 0.00352 |
| cg09385983 | 0.005945 | PLXNA2 | 0.019505 |
| cg13929566 | 0.011286 | SYNE1 | 0.028185 |
| cg10503992 | 0.00069 | TRIM62 | 0.044432 |
| cg04321497 | 0.000464 | TTN | 0.001563 |
| cg13647878 | 0.00897 | TUBA1C | 0.038334 |
| cg02334770 | 0.006588 | UBE2J2 | 0.027877 |
| cg03710176 | 0.006012 | VPS13C | 0.021884 |
| cg01395650 | 0.008254 | ZBTB41 | 0.033364 |
| cg06681008 | 0.008138 | ZNF26 | 0.020245 |

31 differentially methylated probe sites ($p<0.0001$) were associated with 28 differentially expressed genes ($p<0.05$) in human primary astrocytes by control, low glucose, antecedent recurrent low glucose, or recurrent low glucose.
